# Early disruption of the innate-adaptive immune axis *in vivo* after infection with virulent Georgia2007/1 ASFV

**DOI:** 10.1101/2025.06.18.660190

**Authors:** Priscilla YL Tng, Laila Al-Adwani, Lynnette Goatley, Raquel Portugal, Anusyah Rathakrishnan, Christopher L Netherton

## Abstract

Effective immune defence and pathogen clearance requires coordination between innate and adaptive immune responses. However, virulent African swine fever virus (ASFV), which has a high case fatality rate in pigs, causes severe disease by exploiting multiple immune evasion strategies to suppress host responses. The global spread of Georgia2007/1 and its derivatives poses a significant threat to the pig industry and global food security. Although modified live virus vaccines for ASF exist, multiple safety concerns have restricted their use internationally. Conversely, subunit vaccine candidates have not matched the protective efficacy of modified live virus vaccines. This highlights the need to further investigate ASFV-induced immunopathology to support the development of next-generation ASF vaccines. Immune dynamics in whole blood and lymphoid tissues were examined over time after oronasal infection with Georgia 2007/1. CD4^+^ T cells, γδ-TCR^+^ T cells and CD21^+^ B cells were impacted by lymphopenia, and initial immune activation was detected. However, as the disease progressed, impaired maintenance and depletion of adaptive immune cells, such as CD4^+^ T cells and professional antigen-presenting dendritic cells and macrophages was observed. This depletion of cells may have compromised the innate-adaptive immune axis, weakening host ability to mount a robust adaptive immune response and potentially contributing to disease progression. Differential ASFV infection profiles within the spleen were also detected, highlighting the diversity of ASFV cellular tropism. Further investigation into the innate-adaptive immune axis is needed to better understand its role in ASFV infection.

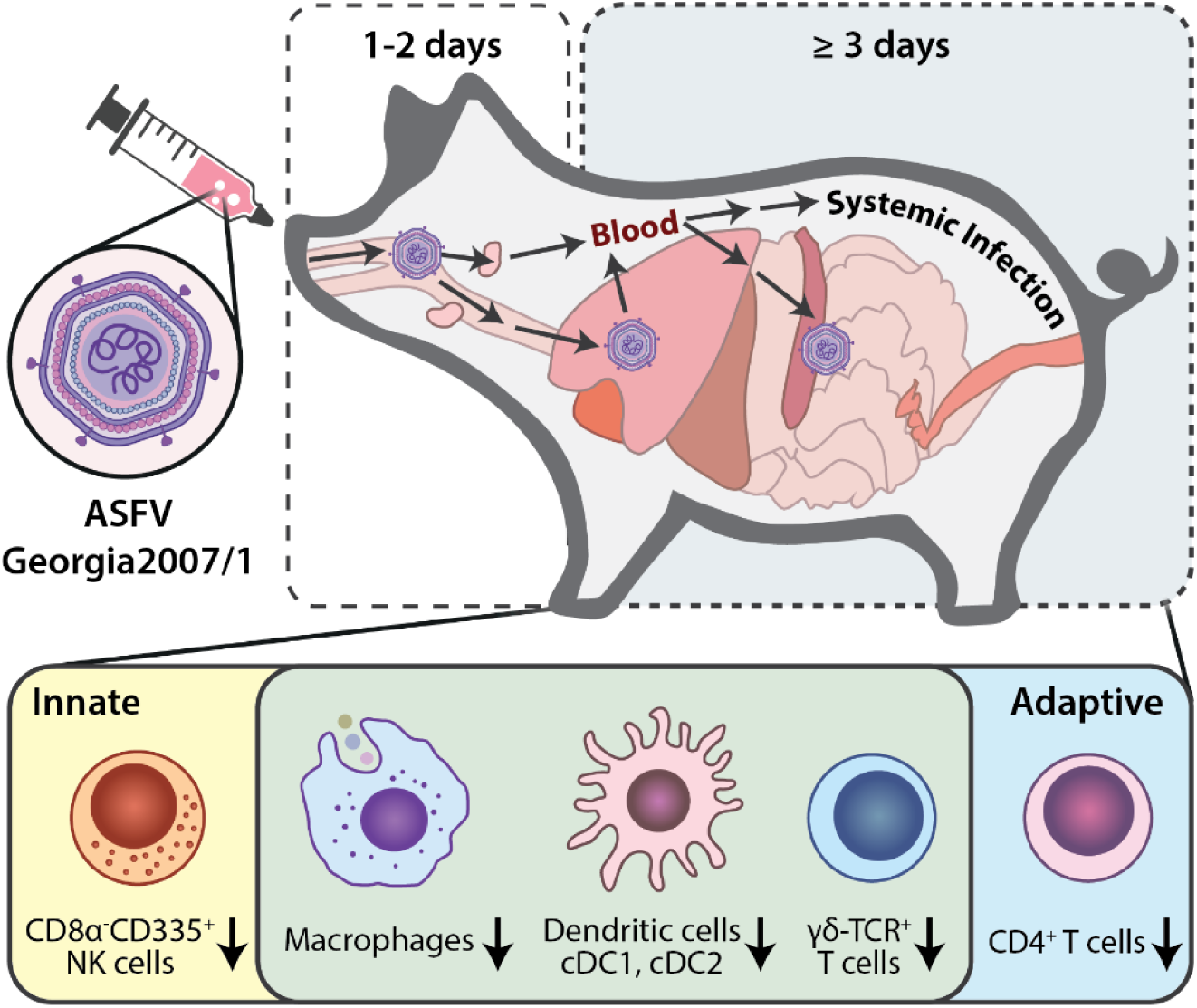

## 2. Introduction

The global pig industry is threatened by the African swine fever (ASF) panzootic. ASF is a notifiable and contagious haemorrhagic viral disease of suids with case fatality rates approaching 100% and is often devastating for affected regions. Since the outbreak of ASF in Georgia in 2007 (isolate Georgia2007/1) [1], ASFV has spread worldwide and outbreaks have been reported on five continents [2]. Based on the current global situation, existing control measures, involving the culling of exposed animals and movement control of animals, are inadequate and have serious socio-economic impacts [3]. Safe and effective ASFV vaccines are urgently needed, but development of such vaccines is challenging due to the complexities of the virus and host immune responses.

Infection with virulent ASFV isolates, such as Georgia2007/1, manifest as an acute disease where pigs suffer high fever, lethargy, and anorexia. Disease progression is rapid: first clinical signs generally manifest between two to seven days post-infection (dpi) and death often within ten days [4]. These events occur before development of adaptive immune responses. Pathological assessments of the course of ASF infection in controlled laboratory settings are often through intramuscular (IM) injection of virus to ensure reliable infection rates with generally synchronised clinical course, however transmission of ASFV in Eurasia is mostly thought to be through close contact with other pigs or ingestion of infectious material. Alternative infection models have been developed to better mimic natural infection routes. One such method is intra-nasal (IN) delivery of virus in pigs with the mucosal atomisation device (MAD) and assessments with influenza demonstrated that this route was able to deliver virus into the lungs [5]. Another method is oronasal inoculation, which has been utilised in more recent work with virulent ASFV isolates [6–8].

To understand the pathogenesis of virulent ASFV, previous work involving sequential sampling of infected animals have focussed on macroscopic lesions, quantification of virus within blood and tissues, virus shedding and predominantly adaptive immune cell function within the blood and selected tissues [6, 9–11]. Other than Greig and Plowright who investigated viral replication at the early sites of infection using ASFV isolates genetically distinct from Georgia2007/1 [10, 11], early virus replication and immune cell dynamics within lymphoid tissues draining initial sites of infection have not been examined in detail for Georgia2007/1. Furthermore, studies on ASFV infection dynamics *in vitro* have primarily focused on monocytes and macrophages within primary macrophage cultures, despite indications that ASFV may target other myeloid cell types [12, 13]. Hence, there is a need for further investigations into the full spectrum of cellular targets of ASFV and their relevance in infection and disease.

Here we first assessed alternative methods of virus inoculation for their reliability and suitability for ASFV challenge experiments. We then sought to unravel the *in vivo* early viral- and immune cell dynamics of both the innate and adaptive immune compartments within secondary lymphoid tissues draining initial infection sites, as well as in visceral lymphoid tissues and blood, after oronasal Georgia2007/1 inoculation. We used highly inbred Babraham pigs to reduce genetic variation in immune responses [14]; immune responses of Babrahams to infection with influenza have been well characterised and demonstrated to be comparable to outbred animals [15]. Furthermore, we previously demonstrated that Babrahams develop acute ASF after infection with virulent ASFV [16]. Besides investigating infection dynamics at early timepoints up to five days post inoculation, we performed multi-parameter spectral flow cytometry (FCM) analysis of both the innate and adaptive immune compartments to provide a more holistic view of the immune landscape within secondary lymphoid tissues and blood following virulent ASFV infection.

## 3. Materials & Methods

### 3.1 Virus

Spleen tissue was obtained from an animal that was infected with the virulent ASFV isolate Georgia2007/1 and was then homogenised in RPMI in a Lysing Matrix A 2 mL tube (MP Biomedicals, USA) with the BeadBug microtube homogeniser at full speed for two minutes. Spleen homogenate was titrated using the haemadsorption assay on porcine bone marrow derived macrophages (PBMs) as previously described [17] and virus titres were calculated as the amount of virus causing haemadsorption in 50% of infected cultures (HAD) with the Spearman-Karber method. Spleen homogenate was diluted to the desired titre with unsupplemented RPMI (Gibco, USA) before oronasal or MAD inoculations. Georgia2007/1 for IM inoculations were grown on PBMs.

### 3.2 Animal Experiments

Female Landrace × large white × Hampshire pigs were sourced from a high health farm in the UK, while both female and male Babraham pigs were bred at the Centre for Dairy Research, University of Reading, Reading, UK. Animals were acclimatised for seven days before any procedures were undertaken. Clinical signs were scored daily, and macroscopic lesions were assessed at post-mortem using previously described methods [18].

#### Experiment 1

Twelve sixteen-weeks-old female Landrace × large white × Hampshire pigs, weighing between 61 – 74 kg, were randomly assigned to three groups for inoculation with Georgia2007/1 using different inoculation methods. AY100 – AY105 (n=6) were oronasally inoculated with 2 × 10^5^ HAD_50_/animal Georgia2007/1 spleen suspension obtained from an infected animal, AZ01 – AZ03 (n=3) were intramuscularly inoculated with 1 × 10^3^ HAD_50_/animal Georgia2007/1 cell culture supernatant and AZ04 – AZ06 (n=3) were intranasally inoculated with 2 × 10^5^ HAD_50_/animal Georgia2007/1 spleen suspension using a mucosal atomization device (MAD, Model: AM501, MedTree, UK). Back-titration of virus inocula revealed virus titres as follows: oronasal inoculation = 5.35 × 10^4^ HAD_50_/animal, intramuscular inoculation = 3.59 × 10^2^ HAD_50_/animal, and intranasal inoculation = 1.08 × 10^5^ HAD_50_/animal. Blinding was not possible during the conduct of the experiment as the virus was administered using different inoculation methods. Whole blood was collected from the animals on −1, 3-, 5-, and 7-days post-inoculation with Georgia2007/1. Tissue samples from the lungs, selected lymph nodes (LN), tonsils and spleen were collected post-mortem for qPCR analysis.

#### Experiment 2

Eleven male (AZ31, AZ32, AZ34, AZ35, AZ36, AZ41, AZ42, AZ43, AZ44, AZ45 and AZ47) and seven female (AZ33, AZ37, AZ38, AZ39, AZ40, AZ46 and AZ48) fifteen- to sixteen-week-old Babraham pigs, weighing between 20 – 37 kg, were used in this experiment. Animal AZ43 arrived with a missing part of the back-leg hoof and was treated with a single application of Terramycin aerosol spray 3.92% and 0.04 mL/kg meloxicam (Metacam) for six days for pain relief. AZ43 and two other animals were randomly assigned to be culled on day 0 of the experiment, and the rest of the animals were oronasally inoculated with spleen suspension of Georgia2007/1 at a titre of 2 × 10^5^ HAD_50_/animal. Back-titration of virus inoculum revealed virus titre to be 2.54 × 10^5^ HAD_50_/animal. Three animals were selected at random to be killed each day from 0- to 5-dpi (n=3 each day). Blinding was not possible during the conduct of the experiment due to the sequential cull experimental design. Whole blood was collected each day from all surviving animals. Post-mortem scoring was performed on all animals immediately after death. Tissue samples were also collected from the lungs, selected LN, tonsils and spleen for qPCR analysis and immuno-phenotyping by flow cytometry (FCM).

### 3.3 Quantitative PCR

ASFV genome copies in whole blood and tissue samples were determined using the assay previously published with slight modifications [19]. Briefly, 20 mg of tissue was homogenised in RPMI with the BeadBug homogeniser as described above. The MagMAX Core nucleic acid extraction kit (Thermo Fisher Scientific, USA) and KingFisher Flex (Thermo Fisher Scientific, USA) were used to extract DNA from homogenised tissue or blood samples, according to the manufacturer’s instructions. qPCRs were performed on a Quantstudio 5 (Thermo Fisher Scientific, USA) with primers from King *et al.* [19]. A two-step thermal profile of 95 °C for 10 minutes and then 45 cycles of 95°C for 15 seconds and 60°C for 60 seconds was used.

### 3.4 Haematological measurements

Fresh EDTA blood samples were subjected to whole blood parameter analysis using the ProCyte Dx Haematology Analyser (IDEXX, USA).

### 3.5 Flow cytometry staining

50 μl of fresh EDTA blood was stained with the following antibodies listed in Table 1 in a final volume of 60 μl at room temperature for 20 minutes. For antibodies labelled with Zenon reagents (Invitrogen, USA), antibodies were conjugated to Zenon labels as per manufacturer’s instructions before staining. Thereafter RBCs were lysed with 450 μl of 1x RBC lysis/fixation solution (Biolegend, USA) for 30 minutes in the dark at room temperature. 250 μl of the cell suspension was acquired on a Cytek Aurora (Cytek Biosciences, USA). Exact cell counts (cells/μl) were determined using the Spectroflo software (Cytek Biosciences, USA).

**Table 1:**
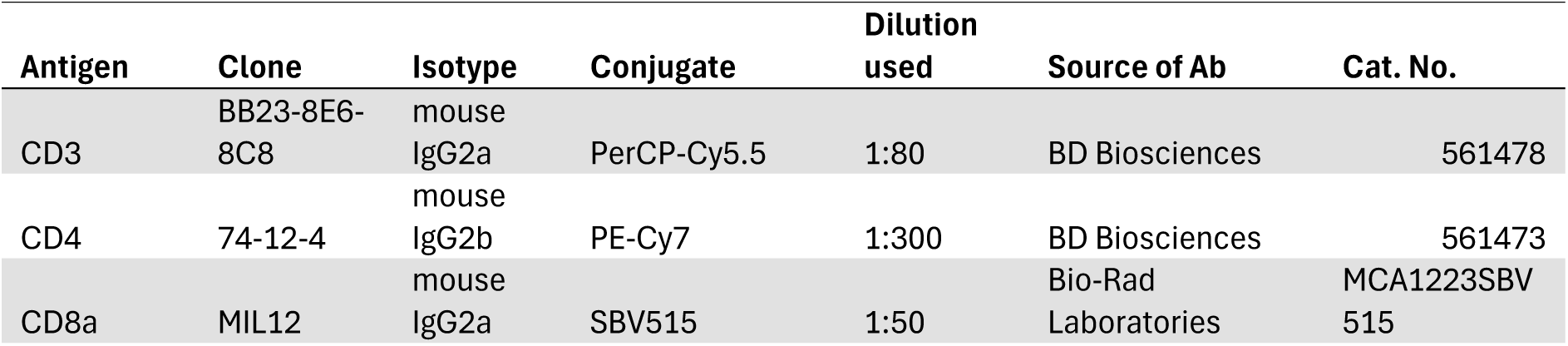
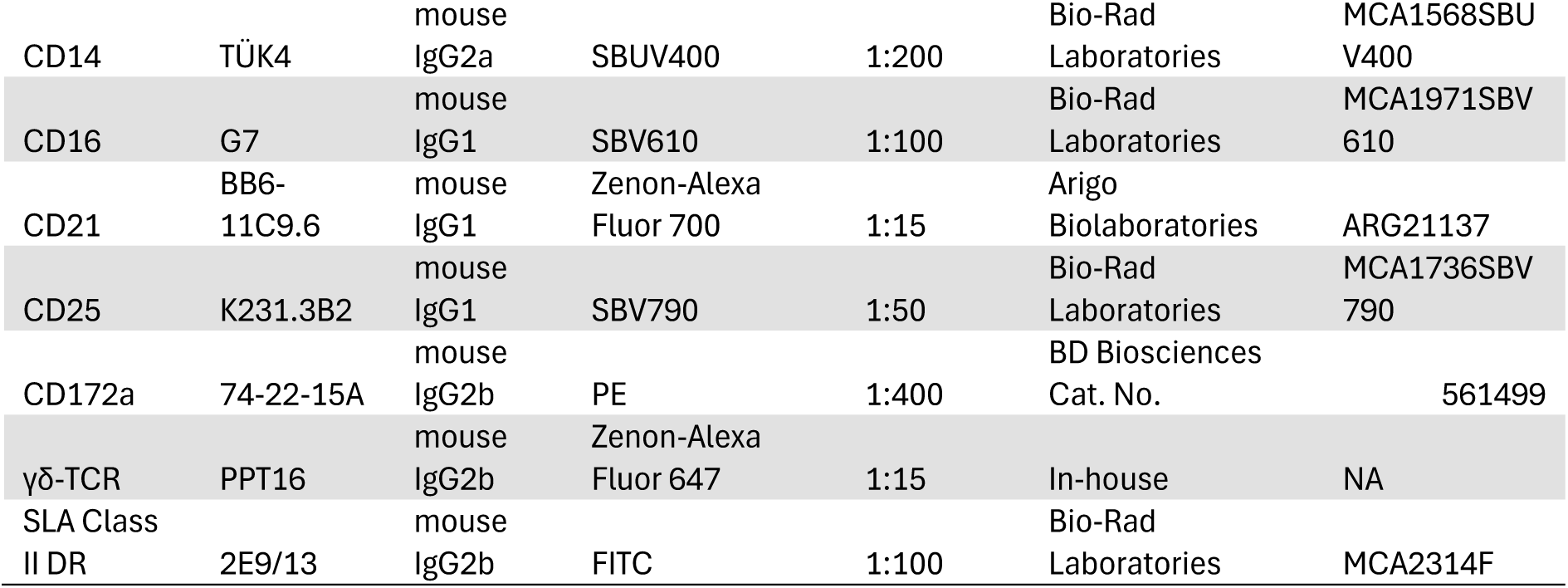
Antibodies used for volumetric whole blood staining.

For lymphocyte and mononuclear phagocyte system (MPS) FCM analyses, selected tissues, spleen, soft-palate tonsil, submandibular-, cervical-, retropharyngeal-, gastro-hepatic lymph nodes were treated with collagenase D (Merck, USA) (at a final concentration of 2.5 mg/ml in unsupplemented RPMI) at 37°C for 30 minutes. Collagenase D was inactivated with by adding EDTA to a final concentration of 10 mM and tissues were mechanically disrupted to obtain single cell suspensions. Cell suspensions were purified further over a histopaque gradient (Merck, USA). Three million cells per tissue were stained immediately after isolation. Cells were first stained with the fixable viability dye eFluor 455UV (Invitrogen, USA, dilution 1:50), before staining with the antibodies outlined in Tables 2 and 3. Where secondary antibodies were used to detect for antibodies binding to extracellular antigens, an additional blocking step with ChromPure Mouse IgG (Jackson Immunoresearch, USA) at a dilution of 1:200 was used before addition of directly conjugated antibodies of the same isotype. Each antibody incubation step was 15 minutes at room temperature and Brilliant Stain buffer Plus (BD Biosciences, USA) was used as per manufacturer’s instructions to prevent polymer-polymer interactions. The cells were fixed and permeabilised with the Foxp3/Transcription Factor Staining kit (Invitrogen, USA) for 30 minutes in the dark at room temperature as per manufacturer’s instructions. Intracellular staining was performed using overnight staining at 4°C. Hybridoma containing antibodies specific for ASFV p72 antigen [20] was conjugated to Zenon mouse IgG2a Alexa-fluor 647 (Invitrogen, USA) before intracellular staining for 15 minutes at room temperature. With both the lymphocyte and MPS FCM panels, at least 60,000 live cells were acquired on the Cytek Aurora (Cytek Biosciences, USA).

**Table 2:** Antibodies used for lymphocyte FCM staining.

| Antigen | Clone | Isotype | Conjugate | Dilution<br>used | Source of Ab | Cat. No. |
| --- | --- | --- | --- | --- | --- | --- |
| *Extracellular* |  |  |  |  |  |  |
| CD2 | MSA4 | mouse<br>IgG2a | NA | 1:200 | Kingfisher | WS0590S-100 |
| mouse IgG2a |  | rat IgG1 | PE-Cy7 | 1:500 | Invitrogen | 25-4210-82 |
| CD3 | BB23-8E6-8C8 | mouse<br>IgG2a | PerCP-Cy5.5 | 1:20 | BD<br>Biosciences | 561478 |
| CD4 | 74-12-4 | mouse<br>IgG2b | Hybridoma | 1:5 | In-house | NA |
| mouse<br>IgG2b | R12-3 | rat IgG2a | BV786 | 1:250 | BD<br>Biosciences | 743179 |
| CD8α | MIL12 | mouse<br>IgG2a | SBV515 | 1:25 | Bio-Rad<br>Laboratories | MCA1223SB<br>V515 |
| CD8β | PPT23 | mouse<br>IgG1 | PE-Cy5 (Lightning-link<br>conjugation) | 1:1600 | Bio-Rad<br>Laboratories | MCA5954G<br>A |
| CD25 | K231.3B2 | mouse<br>IgG1 | SBV610 | 1:50 | Bio-Rad<br>Laboratories | MCA1736SB<br>V610 |
| CD335 | VIV-KM1 | mouse<br>IgG1 | NA | 1:200 | Bio-Rad<br>Laboratories | MCA5972G<br>A |
| mouse<br>IgG1 | A85-1 | rat IgG1 | BUV563 | 1:250 | BD<br>Biosciences | 741254 |
| γδ-TCR | PGBL22A | mouse<br>IgG1 | PE-Texas Red (Lightning-link<br>conjugation) | 1:50 | Kingfisher | WS0621S-100 |
| *Intracellular* |  |  |  |  |  |  |
| CD79a | HM47 | mouse<br>IgG1 | FITC | 1:20 | Biolegend | 986508 |
| FoxP3 | FJK-16s | rat IgG2a | PE-Cy5.5 | 1:400 | Invitrogen | 35-5773-82 |
| Ki67 | b56 | mouse<br>IgG1 | RB780 | 1:2,000 | BD<br>Biosciences | 568762 |
| Perforin | δG9 | mouse<br>IgG2b | Alexa-fluor 647 | 1:50 | BD<br>Biosciences | 563576 |

**Table 3:**
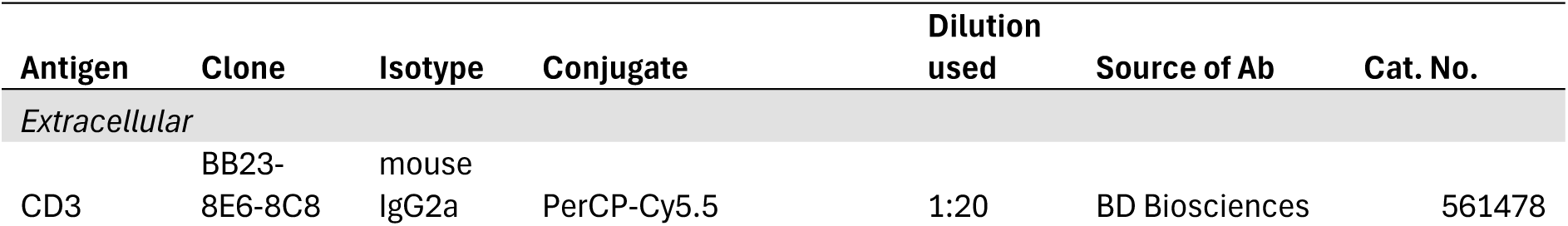
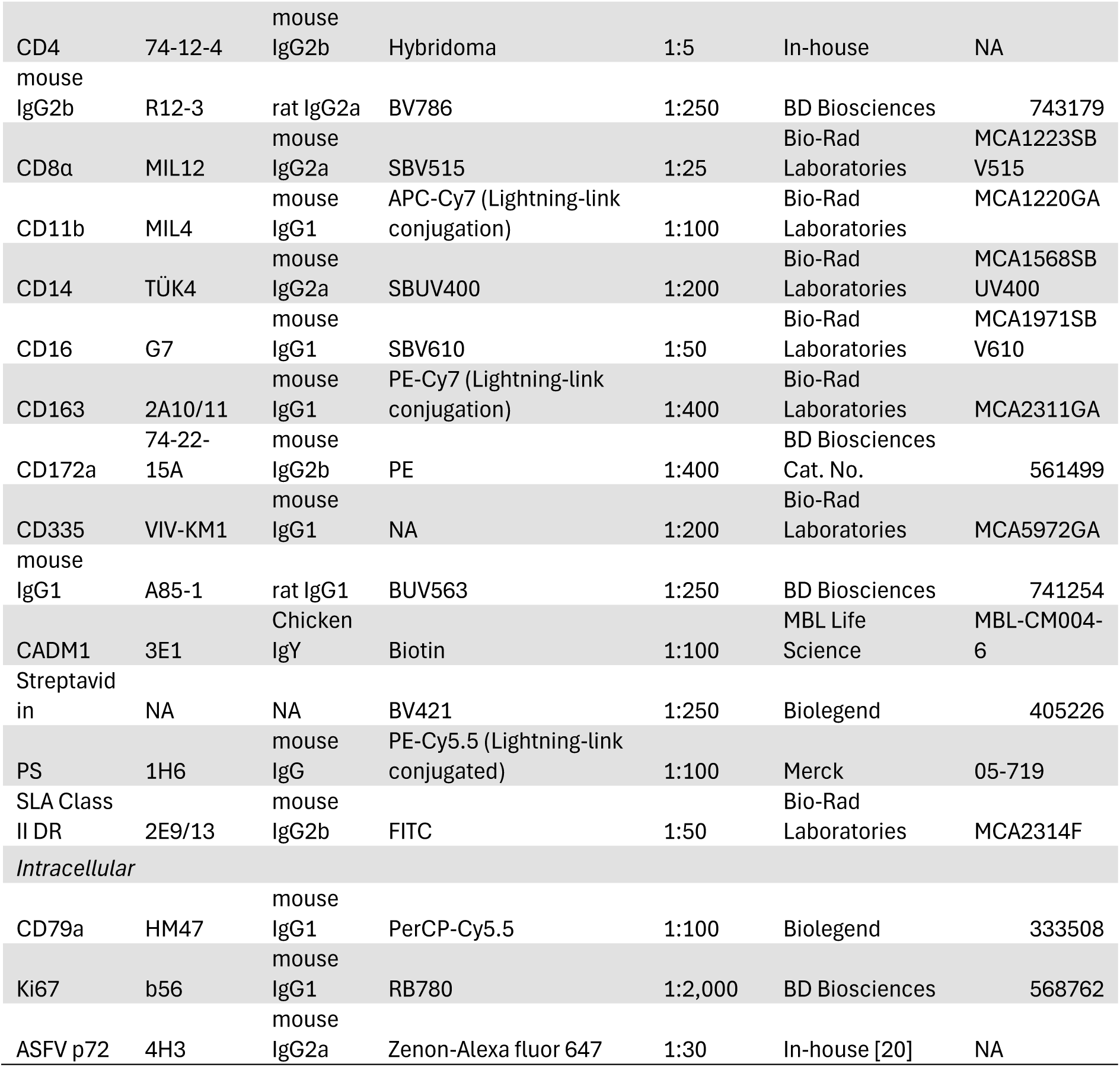
Antibodies used for mononuclear phagocyte system FCM staining.

### 3.6 Flow cytometry analysis

Flow cytometry analysis was performed with FlowJo version 10.10 (BD Biosciences, USA). Citation information for the specific packages used are provided in the Supplementary information. For the analysis of tissue samples, samples were first cleaned with PeacocQC FlowJo package. Specific cell populations (CD3, NK, B cell, myeloid cells) were gated from the live cells and downsampled to obtain equivalent numbers of cells per group. After tSNE unsupervised dimensionality reduction, FlowSOM clustering was performed and clusters were characterised with Cluster Explorer in FlowJo. To avoid over-clustering, only major phenotyping markers for major immune cell populations were included in the analysis due to the limitations of unsupervised dimensionality reduction analysis for rare cell populations [21].

### 3.7 Statistical analysis and visual representation

Statistical analyses were conducted in R (version 4.4.0) and RStudio (version 2023.9.1.494); citation information for the specific packages used are provided in the Supplementary information. One-way ANOVA was preferentially used to investigate the changes in immune cell dynamics and expression of proliferation and cytolytic markers across time, since each animal at each timepoint was an independent event. Data were transformed as appropriate to fit a normal distribution and diagnostic plots of residuals were checked to ensure that there was constant variance between residuals and model assumptions were met. The Kruskal-Wallis test was used where a model could not be fitted or when model assumptions were not met. Tukey’s honest significant differences test (multcomp package) or Dunn’s test (dunn.test package) was used for post hoc analyses where appropriate. Graphs were plotted using the ggplot2 package in R or with GraphPad Prism 10.1.2. Large language models (LLM), ChatGPT versions 3.5 and 4 (OpenAI), were used as tools when generating the code to visualise data in R. LLMs were only used to refine initial R scripts; content was not generated de novo with LLMs. LLM outputs were manually reviewed before sections or none of the outputs were used. Graphs plotted in R were arranged with Illustrator 2024 (Adobe, USA).

## 4 Results

### 4.1 Oronasal infection is a suitable method for ASFV challenge

Here we compared IM to IN and OR inoculation methods (Figure 1A) to establish if these methods were able to result in reliable infection rates in British domestic pigs. Animals started displaying clinical signs such as lethargy and increased temperatures at 4 dpi and some animals within the IN and OR groups showed a delay in the onset of clinical signs in comparison to the IM group (Figure 1B-C, Supplementary Figure 1). Viremia was detected as early as 3 dpi after all methods of inoculation (Figure 1D) and most OR animals reached their humane end points on the same day or one day delayed from the animals in the other groups. Macroscopic scores and viral load in tissues of OR and IN animals were comparable to those in the IM group (Supplementary Figure 2-3). Animals AZ05 (MAD) and AY101 (OR) had to be culled at 7 dpi due to the absence of companion animals within the groups, but these animals were viraemic (Figure 1D), had detectable levels of virus within the tissues assessed (Supplementary Figure 3) and AZ05 was displaying clinical signs typical of ASFV (Figure 1C).

**Figure 1.**
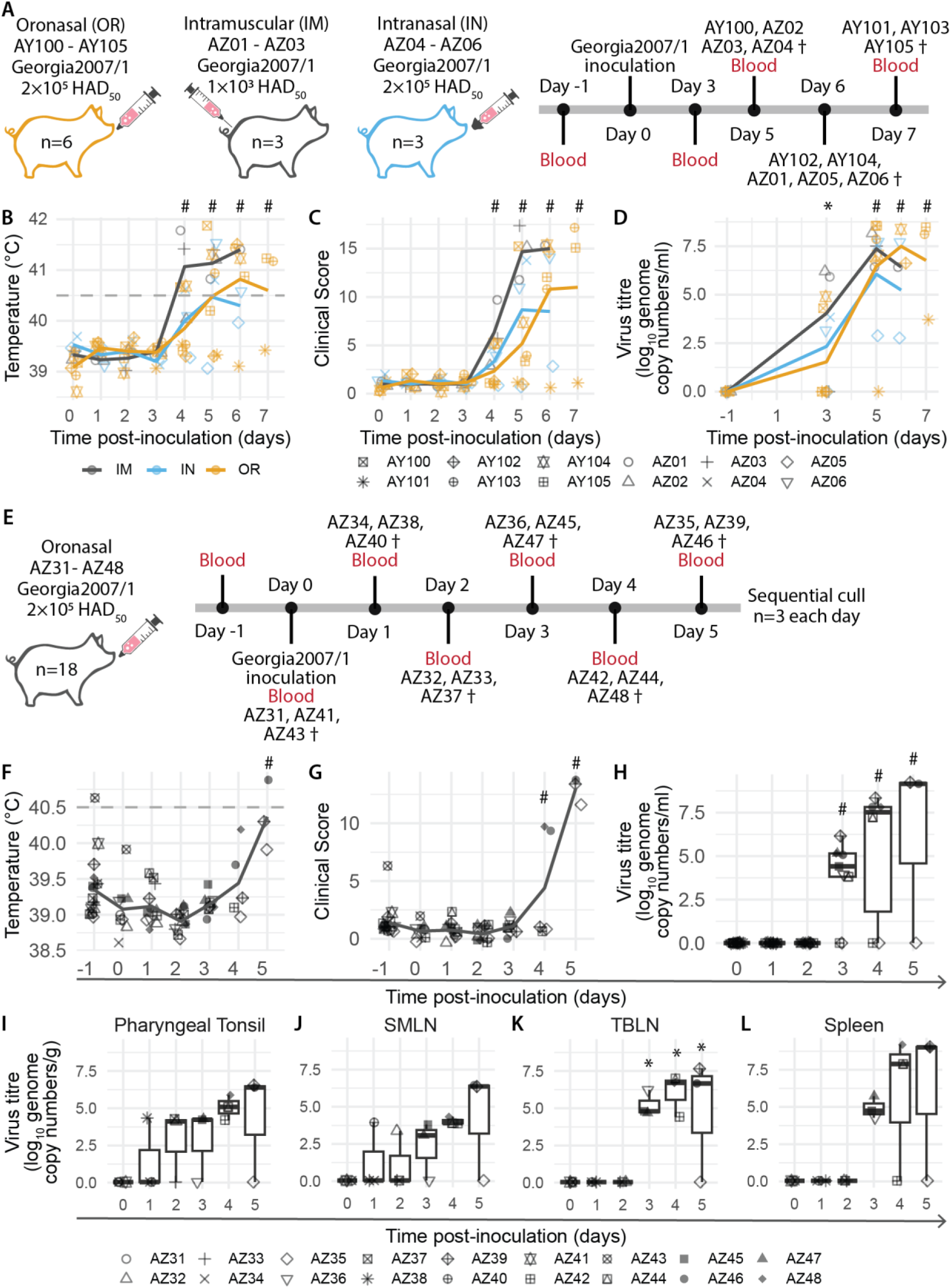
Temperatures and clinical scores after different virus inoculation methods. Temperatures (A) and clinical scores (C) of animals either inoculated through intramuscular injection (IM), intranasally with a MAD device (MAD) or oronasally (OR). The day of virus inoculation is indicated by the arrowhead. Heatmaps of the temperatures (B) and clinical scores (D) of animals in each group. (A, C) Dark grey, yellow and blue lines indicate the mean of each group. (A) Dashed line indicates the temperature at which the animals are considered to have a fever according to the scoring matrix. Each datapoint denotes a single animal. (F-G) Lines depict the mean. (H-L) Median (centreline), the first and third quartiles (box boundaries), maximum and minimum values within 1.5× the interquartile range (whiskers). Statistical significance determined with (B-D) mixed-effects models, (F, H-L) Kruskal-Wallis and Dunn test, and (G) linear model. * p<0.05, # p<0.01.

### 4.2 Early viral replication sites after oronasal inoculation of British domestic pigs

With the confirmation that the pigs could be reliably infected with the oronasal route, we performed a sequential cull experiment with oronasal inoculation of inbred Babraham animals to study the *in vivo* effects and replication sites early in infection with Georgia2007/1. Three animals were culled each day between 0 – 5 dpi (Figure 1E). Similar to the previous experiment, surviving animals started displaying clinical signs at 4 dpi (Figure 1G, Supplementary Figure 4B) and viremia was detected as early as 3 dpi (Figure 1H). Analysis of viral load in draining lymphoid tissues of the infection sites identified that virus could be detected in the pharyngeal tonsil and submandibular LN (SMLN) as early as 1 dpi and levels were sustained between 4 – 5dpi (Figure 1I-J). Within the lungs, virus was detected as early as 1dpi (Supplementary Figure 6H). Virus was found in the tracheobronchial LN (TBLN) and the spleen from 3 dpi and high viral loads were found in the animals that had higher clinical scores (Figure 1K, L). High levels of virus could be detected in the gastrohepatic LN (GHLN) and renal LN (RLN) at 4 dpi (Supplementary Figure 6F, G). AZ35 and AZ42 appeared to have localised infection at the time of cull, but this could be due to a slower course of infection as observed in AY101 in the previous experiment (Figure 1D). AZ35 and AZ42 did not have detectable viremia throughout the study (Figure 1H) but were positive for ASFV in some tissues such as the lungs, TLBN, SMLN and retropharyngeal LN (RPLN) (Figure 1J-K, Supplementary Figure 6E, H). Macroscopic scores for AZ35 at 5 dpi was higher than animals culled at 0 dpi (Supplementary Figure 5). Macroscopic lesions found in the other animals culled on 4 – 5dpi were consistent with ASFV (Supplementary Figure 5) and comparable to those from the previous experiment (Supplementary Figure 2). Taken together, our results indicate that infection begins locally within the draining facial lymph nodes and tonsils, and, to a restricted extent, in the lungs after oronasal inoculation. The infection then progresses to systemic dissemination once the virus is detected in the blood and visceral lymphoid tissues.

### 4.3 Depletion of multiple lymphocyte subpopulations in whole blood after high virulent ASFV infection

Given that virulent ASFV induces lymphopenia [22], we first investigated the changes to whole blood cell populations (Supplementary Figure 7), lymphocyte counts were observed to decrease as the disease progressed in animals that had high viremia (Supplementary Figure 7A). Mean platelet volume (MPV) increased in the sickest animals on 5 dpi (Supplementary Figure 7C), an indication of increased platelet production in response to the infection, and there was a corresponding decrease in reticulocytes (Supplementary Figure 7G), suggesting reduced red blood cell release from the bone marrow of these animals.

Next, using volumetric FCM (Figure 2, Supplementary Figure 8 - 9) we performed more detailed analysis into the affected cell populations. CD3^+^ T cells (Figure 2A) and CD21^+^ B cells (Figure 2H) were observed to decrease at 4-5 dpi, consistent with previous reports of lymphopenia [6, 8, 16]. γδ-TCR^+^CD8α^−^ cells (Figure 2C), CD4^+^CD8α^−^ naïve T cells (Figure 2D), CD4^−^CD8α^+^ T cells (Figure 2E) and CD4^+^CD8α^+^CD25^+^ cells (Figure 2F) contributed to the overall decrease in CD3^+^ T cells. In animals with the most severe disease, CD3^−^CD8α^+^ NK cells decreased over time (Figure 2B) and an increase in CD172a^−^ non-lymphocytes was observed on 5 dpi (Supplementary Figure 9K). In the animals with the most severe clinical disease and viremia (AZ39, AZ46 and AZ48), a transient increase in monocytes was observed on 2 dpi, followed by an apparent downward trend (Figure 2G), similar to IDEXX measurements (Supplementary Figure 7B). These results suggest a dysregulation of whole blood immune cell homeostasis as ASFV infection progresses.

**Figure 2.**
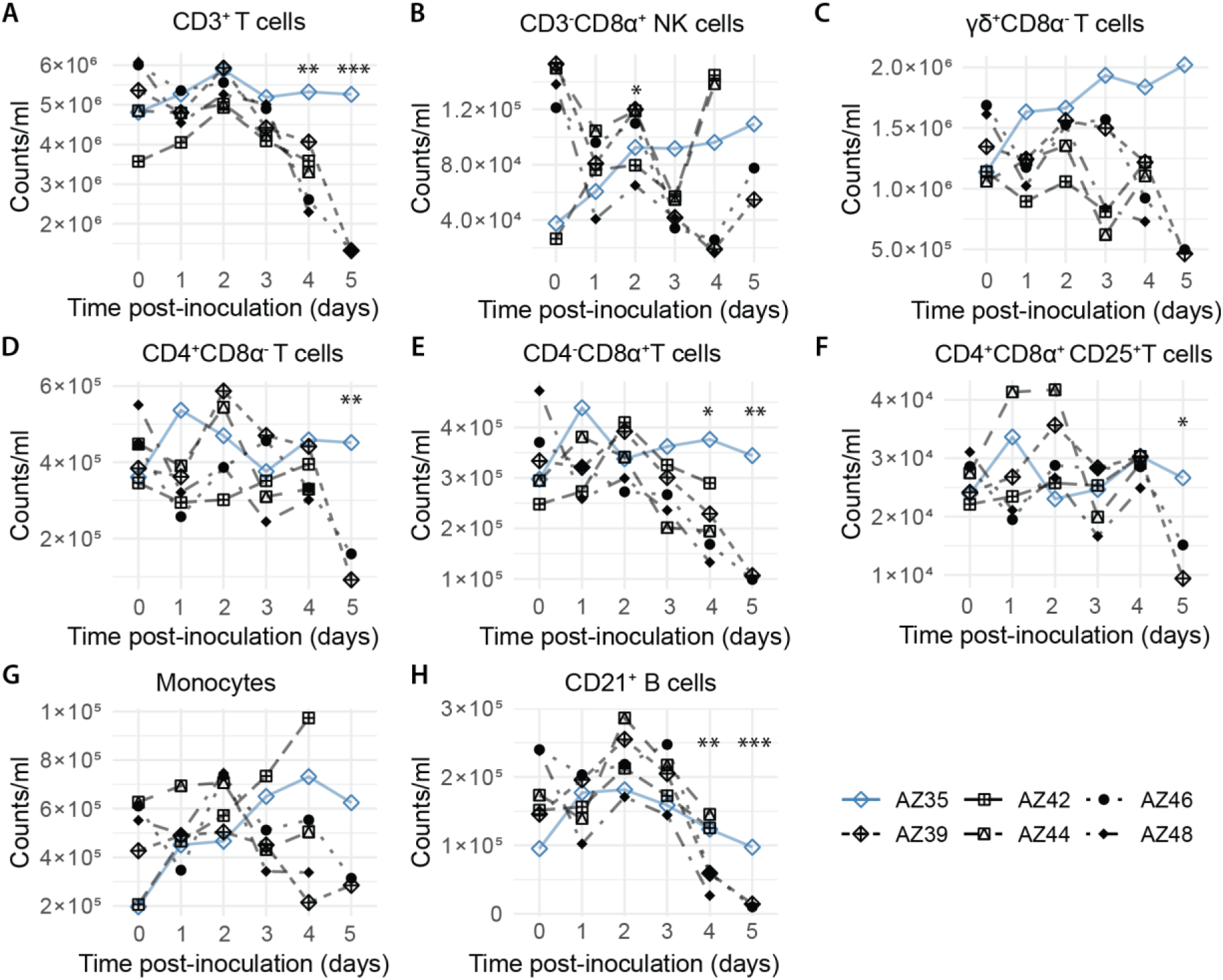
Dynamics of immune cells in whole blood determined with volumetric flow cytometry. Each datapoint denotes a single animal. The animal (AZ35) that was not viraemic on 5 dpi is denoted in blue. Gating strategy can be found in Supplementary Figure 8. Statistical significance determined with mixed-effects model, except for C, L, M and O where linear model was used due to the absence of random effects. ***p<0.001, **p<0.01, *p<0.05.

### 4.4 Dysregulated lymphocyte dynamics within lymphoid tissues during early stages of virulent ASFV infection

Dysfunctional adaptive immune responses after virulent ASFV infection have previously been reported [6, 9]. Here, we sought to examine the dynamics of diverse lymphocyte populations within selected lymphoid tissues during the early stages of ASFV infection. Single cell suspensions derived from the soft palate tonsil (SPTonsil) the LN draining the initial sites of contact (SMLN, CLN, RPLN), as well as the GHLN and spleen, indicators of systemic infection, were subjected to FCM analyses at 0- and 3-5 dpi. We employed tSNE for visualisation, with the caveat that rarer cell types were excluded due to the necessity of downsampling. Thereafter, we performed conventional gating analysis to confirm our tSNE results and to analyse rarer known cell subsets that high unsupervised dimensionality reduction with tSNE could not resolve.

#### γδ-TCR^+^ cells

γδ-TCR^+^ cells are considered rapid responders to infection with multiple protective roles and can be separated into naïve-(CD2^−^CD8α^−^, cluster 5), activated-(CD2^+^CD8α^−^, cluster 4) and effector-(CD2^+^CD8α^+^, cluster 8) γδ-TCR^+^ cells [23, 24]. Since γδ-TCR^+^ cells express cytotoxic markers and perforin is exclusive to CD2^+^γδ-TCR^+^ [25], we also investigated the expression of perforin.

Within the spleen, activated γδ-TCR^+^ levels rose at 3 dpi and remained consistently high throughout the study (Figure 3B, Supplementary Figure 13A). This increase was accompanied by a transient increase in proliferation on 3 dpi and stable levels of perforin expression (Supplementary Figure 13A). Expansion of effector γδ-TCR^+^ was also detected at 3-5 dpi with a corresponding increase in proliferation at 3- and 5 dpi and increased perforin expression 3 dpi (Figure 3B, 4A). A transient increase in activated γδ-TCR^+^ was found at 3 dpi in SPTonsil (Supplementary Figure 13B), while effector γδ-TCR^+^ steadily increased, reaching the peak at 5 dpi (Figure 4B).

**Figure 3.**
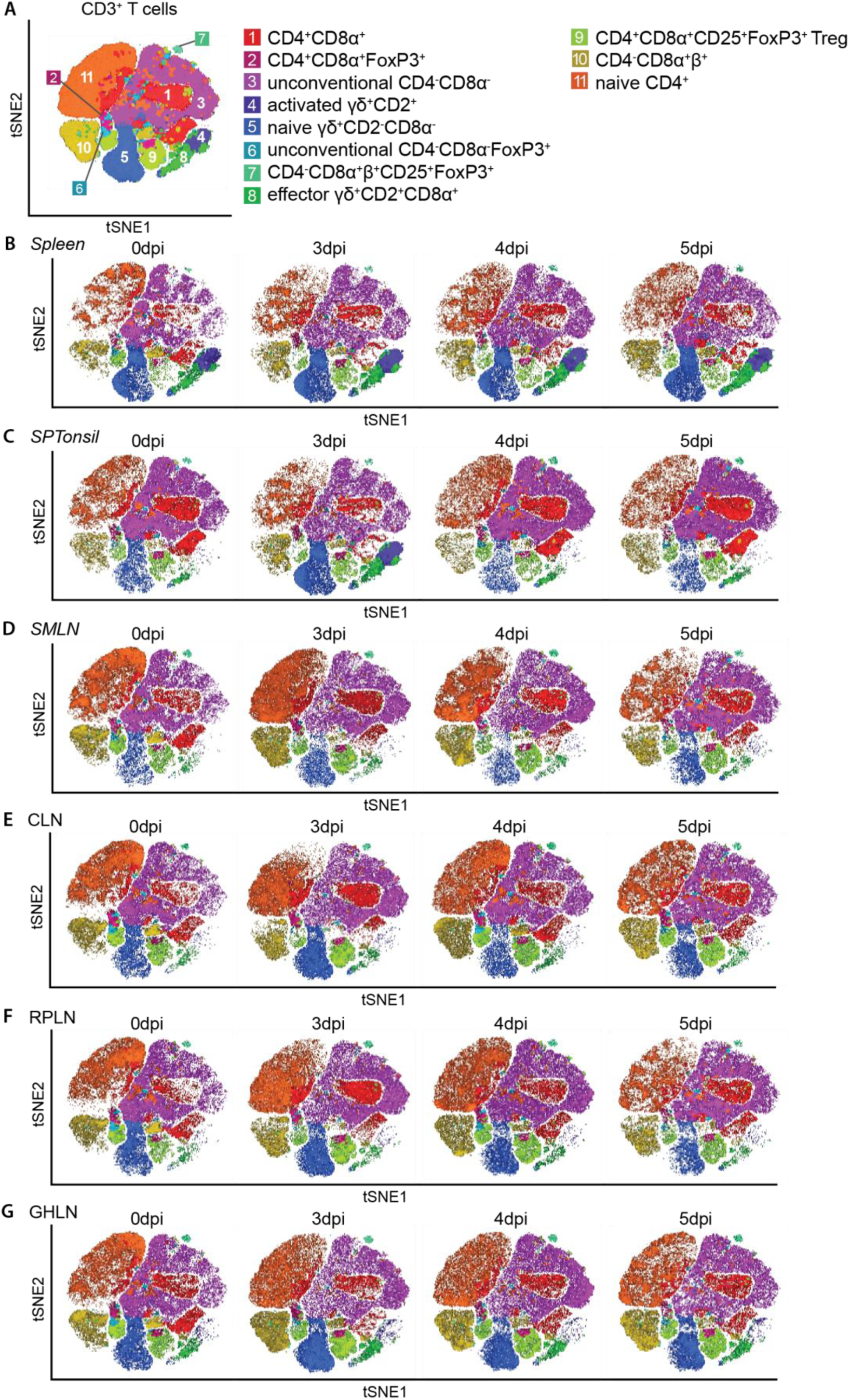
High dimensional analysis of live CD3+ T cells in various lymphoid tissues post-inoculation with Georgia 2007/1. (A) tSNE map overlaid with the eleven clusters obtained with FlowSOM. tSNE maps showing T cell clusters at selected timepoints within the (B) spleen, (C) SMLN, (D) SPTonsil, (E) CLN, (F) RPLN and (G) GHLN. All tissues have n=3 samples at each timepoint except for RPLN 0 dpi (n=2) and 5 dpi (n=1), and GHLN 0 dpi (n=2). CLN: cervical lymph node, GHLN: gastro-hepatic lymph node, RPLN: retropharyngeal lymph node, SMLN: submandibular lymph node, SPTonsil: soft palate tonsil.

**Figure 4.**
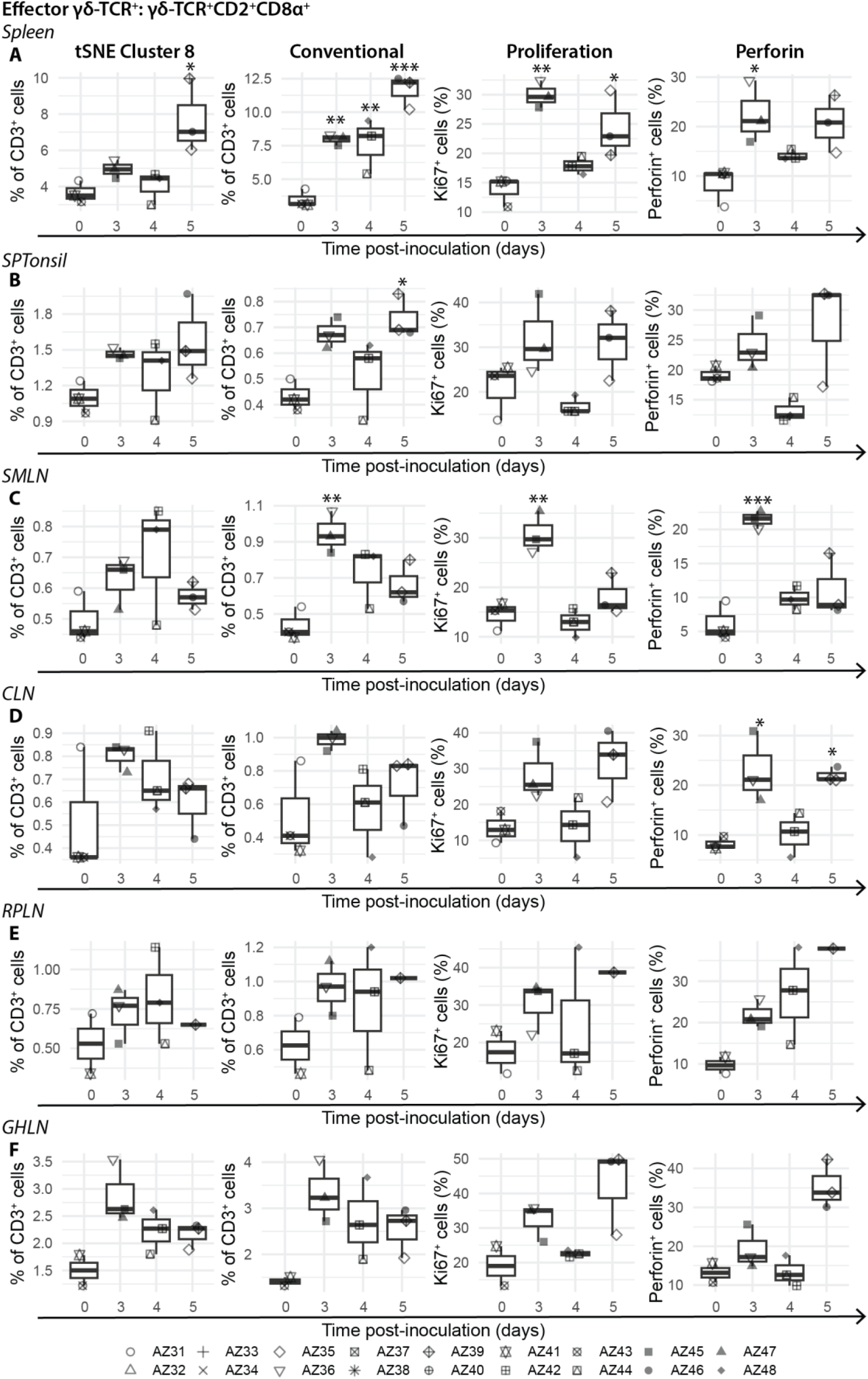
Dynamics, proliferation and perforin expression of effector gdT cells in selected lymphoid tissues. (tSNE cluster 8) high dimensional analysis and (Conventional) conventional gating of activated γδ-TCR^+^ cell dynamics. (Proliferation) Proliferation and (Perforin) perforin expression of activated γδ-TCR^+^ cells. Each datapoint denotes a single animal. Median (centreline), the first and third quartiles (box boundaries), maximum and minimum values within 1.5× the interquartile range (whiskers). Statistical significance determined with one way ANOVA except (M) where Kruskal-Wallis and Dunn test were used. *p<0.05, **p<0.01, ***p<0.001.

In the LNs (Figures 3D-G), brief increases in activated- and effector γδ-TCR^+^ were observed in the SMLN between 3-4 dpi, around the onset of systemic infection (Figure 4C, Supplementary Figures 13C). Similar but less pronounced increases were found in the other LNs assessed (Figure 4D-F and Supplementary Figure 13D-F). Corresponding increases in proliferation and perforin expression in these cell subsets were detected at 3 dpi, especially within the SMLN (Figure 4 and Supplementary Figure 13). However, at 5 dpi where higher viral loads were detected in the LNs (Figure 1J, Supplementary Figure 6D-F), increased proliferation in activated and effector γδ-TCR^+^ did not consistently lead to increased frequencies of these subpopulations (Supplementary Figure 13C-F and Figure 4C-F). Higher perforin expression was detected for effector γδ-TCR^+^ in the SPTonsil and most LNs at 5 dpi (Figure 4).

Similar to activated γδ-TCR^+^, naïve γδ-TCR^+^ briefly increased at 3 dpi within the LNs and had lower levels in the SPTonsil of the sickest animals on 5 dpi, in comparison to the non-viraemic animal, AZ35 (Supplementary Figure 14B-F). Increased proliferation within naïve γδ-TCR^+^ was detected in the spleen and SMLN from 3 dpi, and to a lesser degree in the other tissues assessed (Supplementary Figure 14).

#### Conventional T cells

Following infection with high virulent ASFV, compromised responses from conventional T cells within lymphoid tissues have been observed [6]. To explore this further, we investigated the changes in dynamics within the conventional T cell subsets. Frequencies of double positive (DP) T cells (CD4^+^CD8α^+^β^−^FoxP3^−^, cluster 1) were maintained in most tissues (Figure 3), despite elevated proliferation from 3 dpi (Supplementary Figure 15). In the SMLN, perforin expression was increased in DP T cells at 3-4 dpi but tapered off at 5 dpi (Supplementary Figure 15C). Fluctuations in DP T cell levels across the tissues generally corresponded with inverse fluctuations in naïve CD4^+^ T cells (CD4^+^CD8α^−^β^−^FoxP3^−^, cluster 11) and expansion of naïve CD4^+^ T cells was not observed at 5 dpi despite increased proliferation (Figure 3, Supplementary Figure 16). At 3 dpi, a transient decrease instead of an expansion of cytotoxic T cells (CTLs, CD4^−^ CD8α^+^β^+^FoxP3^−^, cluster 10) was observed in the SPTonsil and to a lesser degree in the facial LN (Figure 3, Supplementary Figure 17), even though proliferation and perforin expression in the CTLs in all tissues assessed were upregulated (Supplementary Figure 17).

Overall, the analysis of major CD3^+^ T cell populations with tSNE was consistent with the results obtained with conventional gating analysis (Figure 3-4, Supplementary Figures 13-17, gating strategy Supplementary Figure 10).

#### T regulatory cells

In contrast to previous reports of upregulation of FoxP3^+^ expressing CD4^+^CD8α^−^ T regulatory cells (Tregs) [6], in our study a rise in DP Tregs (CD4^+^CD8α^+^β^−^CD25^+^FoxP3^+^, cluster 9) instead of the CD4^+^CD8α^−^ Tregs, was observed (Figure 3). Expansion of DP Tregs was evident in the SMLN and SPTonsil, with a similar trend in the spleen, GHLN and RPLN (Supplementary Figure 18). This was associated with increased proliferation in this Treg subset in the spleen and SMLN, and to a lesser extent in the SPTonsil, GHLN and RPLN (Supplementary Figure 19). Similarly, a modest increase in the CD4^−^CD8α^+^β^+^ Tregs (CD4^−^CD8α^+^β^+^CD25^+^FoxP3^+^, cluster 7) was observed in the spleen and LNs at 3-4 dpi, and in the SPTonsil at 5 dpi (Figure 3, Supplementary Figure 18). Increased proliferation in the CD4^−^CD8α^+^β^+^ Tregs was detected at 3 dpi in the spleen and SMLN and a comparable trend was observed in RPLN and GHLN (Supplementary Figure 19).

#### NKT cells

Rarer cell subsets were analysed with conventional gating methods due to the limitations of unsupervised dimensionality reduction analysis [21]. Similar to γδ-TCR^+^ cells, the much rarer invariant NKT are another subset of unconventional T cells that span both innate and adaptive immunity and can be activated either by antigen-dependent or -independent routes [26]. Frequencies of the invariant NKT cell containing population (NKT, CD4^−^CD8α^+^β^−^) declined in the SMLN at 3-4 dpi, while more subtle reduction was observed in the spleen and other LNs (Supplementary Figure 20). Increased proliferation was detected from 3 dpi in the SMLN and to a lesser degree in the other tissues. At the same time, higher perforin expression was found in the spleen and SMLN at 3 dpi, with a similar trend in the other LNs. Conversely, NKT cells transiently increased at 4 dpi, and this was accompanied by reduced perforin levels (Supplementary Figure 20B).

#### NK cells

NK cell subsets with differential expression of CD8α and CD335(NKp56) have been defined previously [27] and CD8α^−^CD335^+^ NK cells were described to be highly activated with high cytokine expression and cytolytic activity [28]. A modest increase in CD8α^−^ CD335^+^ NK was observed in all tissues at 3 dpi, which corresponded to proliferation levels detected, but this rise was largely reversed to below 0 dpi levels by 5 dpi (Supplementary Figure 21). Increased perforin expression was observed in splenic CD8α^−^CD335^+^ NK cells from 3 dpi, and much later at 5 dpi in the LNs of animals with more severe disease (Supplementary Figure 21). Perforin expression in this NK cell subset was reduced from 3 dpi in the SPTonsil and did not recover (Supplementary Figure 21B). CD8α^+^CD335^−^ NK cells were observed to decline all tissues at 3 dpi (Supplementary Figure 22), accompanied by a rise in proliferation and perforin levels. Correspondingly, expansion of CD8α^+^CD335^+^ NK levels, together with increased proliferation, was found at the same time (3 dpi) in LNs and SPTonsil (Supplementary Figure 23), which may be attributed to the upregulation of CD335 expression in CD8α^+^CD335^−^ NK cells [27].

#### B cells

Proliferation was evident in the B cells in SMLN from 3 dpi with a trend towards increased B cell frequencies at 5 dpi (Supplementary Figure 24C). Similar increase in B cell levels were observed in the other facial LNs. This increase could be due to the decrease in CD3^+^ T cells in these LNs (Supplementary Figure 12C-E). Conversely, while B cell proliferation was detected in the spleen at 3 dpi, this was not accompanied by a rise in B cell frequencies (Supplementary Figure 24A), even though overall CD3^+^ T cell frequencies were stable at this time point (Supplementary Figure 12A). Increased proliferation may have contributed to the rise in B cell frequencies in SPTonsil at 5 dpi (Supplementary Figure 24B) due to maintained levels of CD3^+^ T cells (Supplementary Figure 12B).

Taken together, the alterations of T-, NK- and B cell compartments in the early stages of virulent ASFV infection suggest a disruption in the effector cell populations of the adaptive immune system that is characterised by a failure to maintain key immune cell populations and inconsistent proliferative responses.

### 4.5 Disruption and depletion of mononuclear phagocyte system (MPS) cells involved in innate-adaptive immune cross talk following virulent ASFV infection

Here we investigated the impact of infection with virulent ASFV on innate immune cells from the MPS since these are equally as important as mediators of adaptive immunity, especially in the primary immune response. Single cell suspensions subjected to FCM analyses were stained with a separate panel focusing on the MPS. Similar to lymphocyte analyses, samples from tissues were visualised with tSNE (Figure 5) and professional antigen presenting cell (APCs) were subjected to conventional gating analysis to analyse rare cell subsets (Supplementary Figures 27-29). In general, unsupervised dimensionality reduction analyses across the tissues aligned with conventional gating analyses except for rarer populations.

**Figure 5.**
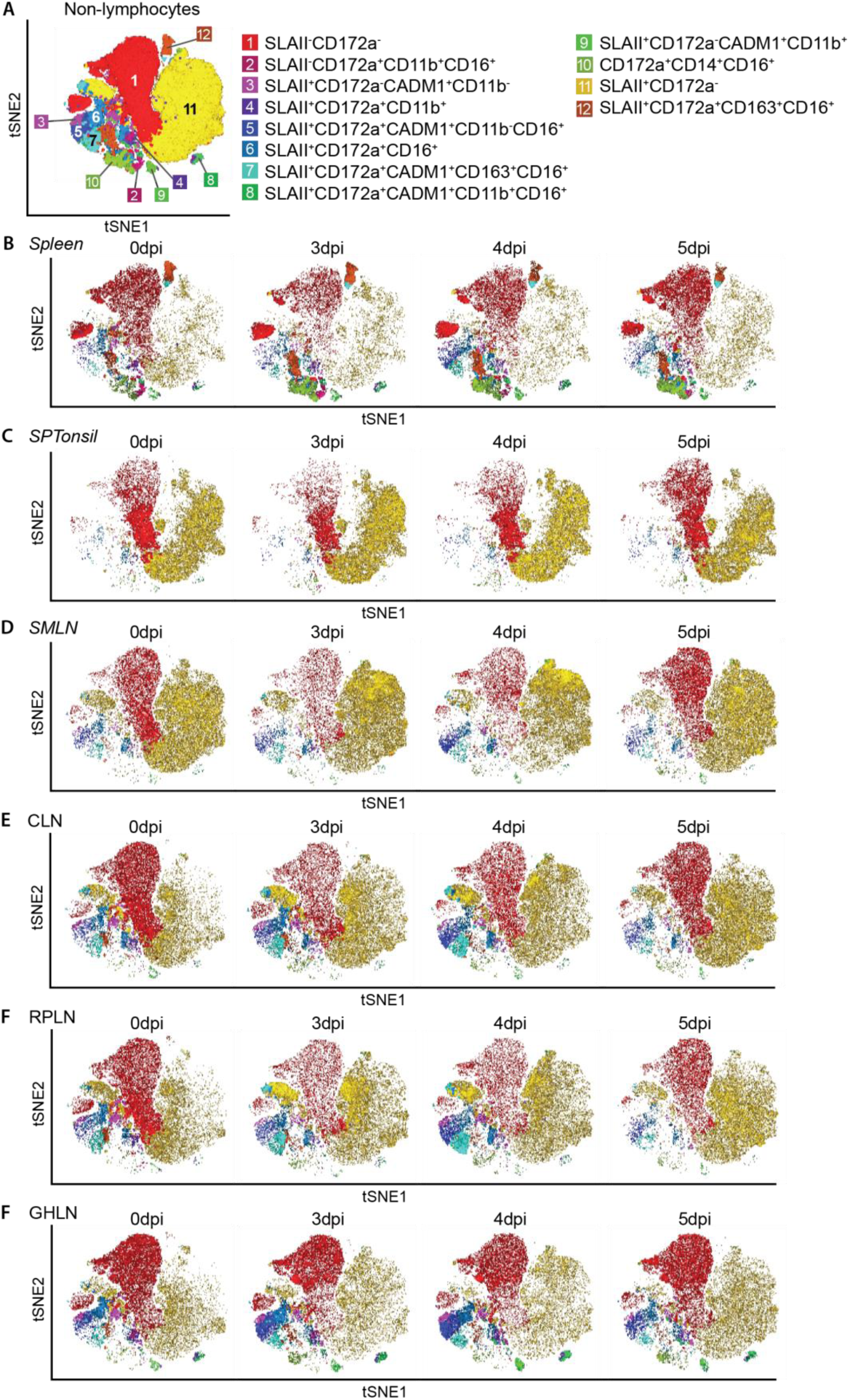
High dimensional analysis of live non-lymphocyte in various lymphoid tissues post-inoculation with Georgia 2007/1. (A) tSNE map overlaid with the twelve clusters obtained with FlowSOM. tSNE maps showing non-lymphocyte clusters at selected timepoints within the ((B) spleen, (C) SMLN, (D) SPTonsil, (E) CLN, (F) RPLN and (G) GHLN. All tissues have n=3 samples at each timepoint except for RPLN 0 dpi (n=2) and 5 dpi (n=1), and GHLN 0 dpi (n=2). CLN: cervical lymph node, GHLN: gastro-hepatic lymph node, RPLN: retropharyngeal lymph node, SMLN: submandibular lymph node, SPTonsil: soft palate tonsil.

#### Dendritic cells

Frequencies of antigen presenting non-lymphocytes expressing SLA class II DR (SLAII) increased in the spleen between 3-4 dpi (Supplementary Figure 27A) and a similar expansion, but to a lesser extent, were observed in SPTonsil, CLN and RPLN 3 dpi (Supplementary Figure 27B, D-E). This increase was largely reversed by 5 dpi. Dendritic cells are key APCs modulating both innate and adaptive immunity [29] and porcine conventional dendritic cells (cDC) have previously been defined as CD14^−^CD172a^−/lo^CADM1^+^CD11b^+^ for cDC1 (Figure 5 cluster 9) and CD14^−^CD172a^+^CADM1^+^CD11b^+^ for cDC2 (Figure 5 cluster 8) [30]. Depletion of putative cDC1 cells was observed in all tissues from 3 dpi (Figure 6A, Supplementary Figure 28), especially in the SPTonsil and CLN, even though these cells are known to migrate to LNs and proliferate in response to infection [29]. Likewise, reduced levels of putative cDC2 were detected in all tissues by 5 dpi (Figure 6C, Supplementary Figure 29), although a transient increase in putative cDC2 was evident in the spleen at 4 dpi (Figure 6C). Increased proliferation was detected in both cDC1 and cDC2 populations in the spleen from 3 dpi (Figure 6A, C), but this was insufficient to boost frequencies in either subset. Levels of both cDC1 and cDC2 subsets were severely depleted in the animals with the most severe clinical signs at 5 dpi (Figure 6A, C, Supplementary Figure 28-29).

**Figure 6.**
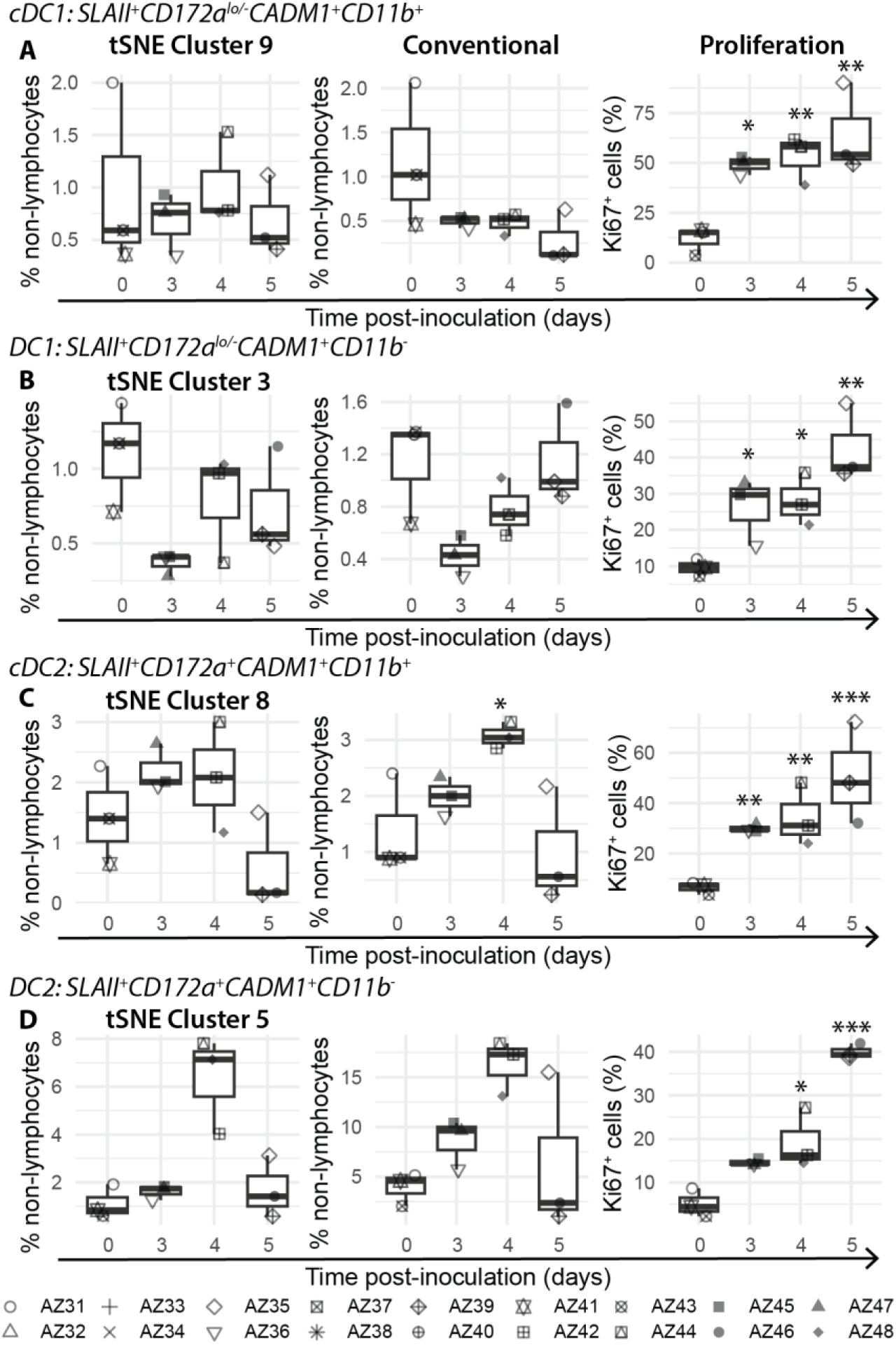
Dynamics and proliferation of dendritic cell population in the spleen after inoculation with Georgia 2007/1. (tSNE cluster) high dimensional analysis and (Conventional) manual gating of dendritic cell dynamics within the spleen. (Proliferation) Proliferation of spleen derived dendritic cell subsets. Each datapoint denotes a single animal. Median (centreline), the first and third quartiles (box boundaries), maximum and minimum values within 1.5× the interquartile range (whiskers). Statistical significance determined with one way ANOVA. *p<0.05, **p<0.01, ***p<0.001.

Other CADM1^+^ cell subsets were also affected as disease progressed. Here, we define SLAII^+^CD172a^lo/-^CADM1^+^CD11b^−^ cells as DC1 and SLAII^+^CD172a^+^CADM1^+^CD11b^−^ cells as DC2. Within the spleen, DC1 frequencies briefly decreased at 3 dpi, and this was accompanied by increased proliferation that was maintained thereafter (Figure 6B). Conversely, a transient increase in DC2 within the spleen was observed at 4 dpi, despite increased proliferation from 4 dpi (Figure 6D). A transient increase in DC1and DC2 levels was evident within facial LNs, and DC2 within GHLN, at 4 dpi (Supplementary Figures 28-29).

#### Monocytes and macrophages

Monocytes and macrophages play central roles in the MPS and are key replication sites for ASFV [31]. Monocyte (tSNE Cluster 10, CD172a^+^CD14^++^) frequencies rose in the spleen and SMLN between 3-5 dpi (Figure 5B, D) and to a lesser degree in the SPTonsil and CLN at 3 dpi (Figure 5C, E), coinciding with the reduction in circulating monocytes in the blood of the most viraemic animals (Figure 2G). More detailed analysis of SLAII^+^ monocytes (SLAII^+^CD172a^+^CD14^++^) found a peak increase in monocytes within the spleen and SMLN at 4 dpi but levels were on a downward trend at 5 dpi (Supplementary Figure 30A, 31B). Similar SLAII^+^ monocyte dynamics were observed in SPTonsil, CLN and RPLN (Supplementary Figure 31A, C-D). As expected, CD163+ macrophages (tSNE Cluster 12, SLAII^+^CD172a^+^CADM1^−^CD163^+^) were depleted by 5 dpi in all tissues and was most severe in the spleens of animals with the highest viremia on 5 dpi (Figure 5, Supplementary Figure 30B, 31). In stark contrast to the depletion of monocytes and macrophages, proliferation of splenic monocytes and macrophages was upregulated at 3 dpi and peaked at 5 dpi (Supplementary Figure 30).

Monocytes are known to replenish macrophages in tissues during inflammation and infection [32], and monocyte derived macrophages (mo-macrophages) within the porcine lung have been previously defined as SLAII^++^CD11b^+^CD14^+^CD163^int^CADM1^lo^ cells [33]. We identified SLAII^++^CD11b^+^CD14^+^CD163^+^ (defined here as mo-macrophages) in the spleen and subdivided these into CADM1^+^ (CADM1^+^ mo-macrophages) and CADM1^−^ (CADM1^−^ mo-macrophages) subsets (Supplementary Figures 32-33). Frequencies of mo-macrophages increased between 3-4 dpi but were depleted in the spleens with the highest viral loads (AZ39, AZ46, AZ48) (Supplementary Figure 33A). The CADM1-subset of mo-macrophages was elevated earlier at 3 dpi than the CADM1+ subset at 4 dpi (Supplementary Figure 33B-C).

#### Apoptosis

We detected haemorrhagic lymphadenitis within sites of early replication at 5 dpi (Supplementary Figure 5) and depletion was evident in some of the major cell populations (Supplementary Figure 12, 27). Hence, we sought to determine if apoptosis was a contributing factor in lymphoid tissues draining the initial sites of replication. Using the fixed apoptosis necrosis assay [34], we determined that early apoptosis was evident in the SMLN, SPTonsil and CLN at 4 dpi and peak levels of early apoptosis in the lymphocyte compartment containing both CD3^+^ T cells and CD79a^+^ B cells was at this timepoint (Supplementary Figure 35). Slightly different trends were observed for antigen presenting non-lymphocytes in these tissues. Steady increase in early apoptosis was observed in the SPTonsil between 3-5 dpi (Supplementary Figure 35A), while apoptosis levels fluctuated around the baseline in the SMLN and CLN (Supplementary Figure 35B, C).

#### ASFV positive cells within the MPS

Lastly, we used an antibody specific for the ASFV p72/B646L protein to phenotype infected cells in the spleens of animals with the highest virus titres (AZ39, AZ46 and AZ48, Supplementary Figures 36-37)[20]. Although the titres detected were between 10^9.02 – 9.19^ genome copies/g spleen, we were only able to detect 0.82 ± 0.51% (mean, SD) of live cells that were infected. The major population of cells infected were non-lymphocytes of the MPS (Supplementary Figure 36A). AZ48 displayed clinical signs of ASF at 4 dpi and of the non-lymphocyte cells identified to be p72^+^, antigen presenting CD172a^+^CD14^−^CADM1^+^CD11b^−^ cells were a major subpopulation (Supplementary Figure 36B). In AZ39 and AZ46, the top three p72^+^ non-lymphocyte cells were non-antigen presenting SLAII^−^CD14^−^CD172a^+^, non-antigen presenting monocytes (SLAII^−^ CD14^++^) and antigen presenting monocytes (SLAII^+^CD14^++^) (Supplementary Figure 36B). Of the immunophenotyped cell subsets within the spleens, macrophages (CD172a^+^CD11b^−^CADM1^−^CD163^+^) had the highest susceptibility, followed by a population of non-antigen presenting SLAII^−^CD14^−^CD172a^+^ cells.

Collectively, these observed dynamics from the MPS indicate that there is an attempt to mount a primary response upon ASFV infection. However, the depletion of professional APCs, the inability to sustain the replacement of these cells, and increased apoptosis of APCs in lymphoid tissues likely contribute to a dysfunctional response.

## 5 Discussion

In this study we sought to characterise the *in vivo* dynamics of primary responses in the blood and lymphoid tissues at the early stages of ASFV infection. We found that despite an initial attempt to mount an immune response, clinical disease was accompanied by a dysregulated maintenance and depletion of immune cell populations, such as the CD4^+^ T cells, γδ-TCR^+^ T cells and the cDCs, essential for the primary immune response and for bridging adaptive and innate immune responses.

We first validated the OR and IN infection methods with the conventional IM injection infection route in outbred pigs and both routes were able to induce reliable infection, albeit with a noticeable delay in onset of clinical signs and viremia in some of the animals. Although infections in OR and IN groups were not as synchronised as the IM animals, all animals were infected with ASFV, as detected by qPCR. Previous work into the delivery of virus intranasally with the MAD device demonstrated that this method delivered a high proportion of the virus into a restricted area of the lungs [5]. Our back titrations demonstrated that the animals could be reliably inoculated with lower virus titres derived from infected spleen than previous work with Armenia2008 [6], which is effectively identical to Georgia2007/1.

Next, we investigated the primary responses and immune cell dynamics in early stages of ASFV infection in inbred Babraham pigs using the OR infection route. Immune responses of Babraham pigs after H1N1 influenza infection have previously been found to be comparable to that of outbred animals [15] and have been used in ASF infection studies [16]. ASF typical clinical signs and macroscopic lesions were observed in animals culled at 4-5 dpi, but qPCR analyses demonstrated that AZ35 and AZ42 had not develop systemic infections at the point they were killed. It is possible that these animals had a delayed infection course and would have developed systemic infections as demonstrated by AY101 in the first experiment at 7 dpi. These results highlight the need for diagnostic qPCR detection of ASFV to determine infection since some of the clinical signs and macroscopic lesions typical of ASFV infection are similar to that of other porcine diseases.

Early detection of ASFV in the pharyngeal tonsils and SMLN in this study were consistent with previous results after intranasal inoculation of the Tengani isolates [11]. In addition, virus was detected in the lungs of two animals (AZ38 and AZ40) culled on 1 dpi. It is possible that these animals inhaled the virus deeper into the lungs during the inoculation as animals were only restrained and not sedated for virus inoculation, which may be similar to intranasal delivery with a MAD device [5]. Detection of virus in the blood, spleen and other visceral lymph nodes from 3 dpi were indicative of virus dissemination leading to systemic infection and it appears that virus is reintroduced to the facial lymphoid tissues during this phase.

Since systemic infection typically manifests from 3 dpi, we characterised the dynamics of major immune populations of the blood and lymphoid organs that contribute to the primary response post-infection. Consistent with findings from virulent Armenia2008 and CADC_HN09 infection of outbred pigs [6, 9], we observed lymphopenia, which could be attributed to the loss of CD3^+^CD4^+^CD8α^−^ and CD3^+^CD4^+^CD8α^−^ T-and CD21^+^ B cells. Given that immune cells migrate from the bloodstream to lymphoid tissues during infection [35], we investigated whether immune cell migration could contribute to the observed lymphopenia. In general, CD3^+^ T cell frequencies were decreased while B cell frequencies were maintained across the tissues by 5 dpi, indicating that the loss was not due to migration. This is similar to findings from single cell RNA sequencing (scRNAseq) of the spleens of infected animals [12]. Literature suggests that apoptosis of lymphocytes may be a contributing factor [36]; similarly, we detected higher levels of early apoptosis in the lymphocyte population (comprising of CD3^+^ T- and CD79a^+^ B cells) from 4 dpi in the facial lymph nodes. These findings lead us to speculate that the loss in circulating CD3^+^ T cells may predominantly be due to apoptosis, as previously shown [37], while loss of B cells may delayed and driven by the depletion of T-helper cells [36, 37]. It is possible that the drastic B cell depletion in tissues observed in other studies was not detected here due to the earlier study termination at 5 dpi. A more detailed analysis into B cell dynamics and B-cell specific apoptosis is necessary to confirm these findings, particularly due to the different B cell antibodies used in whole blood (CD21) and tissue (CD79a) FCM analyses and the inability to differentiate B cells from T cells when assessing apoptosis.

Another important subset of CD3^+^ T cells that are impacted by virulent ASFV are γδ-TCR^+^ cells that are known to be cytolytic and are involved in innate-adaptive crosstalk [38]. Similar to findings from Armenia2008 infections [6], we observed a reduction in γδ-TCR^+^ cells in the blood of animals with more advanced clinical disease, indicative of a dysregulated maintenance of these cells. However, unlike the Armenia2008 results, we detected a transient increase in perforin expression in effector γδ-TCR^+^ cells at 3 dpi across the tissues. This observation aligns with in vitro data, where resting γδ-TCR^+^ cells exhibited little to no perforin expression, but upregulated perforin along with an increase in effector γδ-TCR^+^ cells after co-culture and activation with macrophages [25], which may be a contributing factor at 3 dpi. The fluctuating levels of perforin in γδ-TCR^+^ cells between 4-5 dpi may reflect perforin consumption, potentially through cytotoxic activity, though this requires confirmation in future studies alongside other killing markers such as Fas/FasL [6, 25]. Besides cytotoxic activity, γδ-TCR^+^ have been implicated in innate-adaptive crosstalk, including the priming and activation of other immune cells such as DCs and NK cells to boost adaptive immune responses [38–40]. Furthermore, reciprocal enhancement and complementary function between γδ-TCR^+^ and DCs have been reported [41, 42], but this remains to be explored further in pigs, which have significantly more γδ-TCR^+^ cells than mice and humans.

In contrast to the increase in iNKT after Armenia2008 infection [6], we detected overall reductions in this population. Notably, our staining only identified the CD4-CD8α+ subpopulation of iNKT that are considered to be naïve iNKT due to the absence of staining with the CD1d tetramer. Previous work with Armenia2008 did not measure iNKT levels in the spleen or facial LN, and no decrease in splenic iNKT was observed with moderately virulent ASFV, Estonia2014. Therefore, further studies are needed to compare the effects of ASFV isolates with varying virulence on iNKT populations.

NK cells are another critical subset of immune cells that bridge the innate-adaptive axis. From our results, we speculate that highly activated cytolytic CD8α^−^CD335^+^ NK cells were depleted across the tissues, and these were replaced by CD8α^+^CD335^+^ NK cells. It has been shown that CD8α^+^CD335^+^ NK cells have lower cytokine expression, degranulation and cytolytic capabilities in comparison to CD8α^−^CD335^+^ NK cells [28]. It is possible that the loss of CD8α^−^CD335^+^ NK cells contributed to a dysfunctional immune response and disease severity.

Tregs, which have been described to contribute to the control of tissue damage in infection [43], were also investigated in infections with Armenia2008, where CD4+CD8α^−^ Tregs were upregulated at 7dpi in the blood, spleen and GHLN [6]. Conversely, a decrease in blood CD4+ Tregs, which includes both CD4+CD8α^+^ and CD4+CD8α^−^ Tregs, was found in infections with CADC_HN09. Our study had a shorter duration than these studies, but similar to Tian *et al*.[9] we observed reductions in the CD4+ Treg containing populations of the blood (CD4+CD8α^−^CD25^+^ and CD4+CD8α^−^CD25^+^T cells). Transient increase in frequencies and proliferation in CD8α^+^ Tregs detected in the tissues we examined could be an attempt to regulate cellular responses after infection as identified in human lymphoid tissues [44], but as suggested previously, the effects of Treg dysregulation in acute ASFV infection require more in-depth exploration.

Apoptosis could have contributed to the depletion of antigen presenting components of the MPS in the facial LNs. T cell responses are crucial for protection against ASFV, and impaired T cell responses have been described after virulent ASFV infection [6, 9, 45]. The depletion of SLAII^+^ APCs by 5 dpi in the animals with more advanced disease may contribute to impaired development of an acquired immune response due to reduced activation of T cells [46].

cDCs are primary APCs that bridge the innate and adaptive immune responses as key mediators of the T cell response, and these are typically recruited into the draining LNs after infection [29]. Across the secondary lymphoid tissues assessed, putative cDC1 and cDC2 populations were generally depleted, especially in animals with the highest viral loads, despite increased proliferation. Interestingly, there was a transient increase in putative cDC2 4 dpi in the spleen. Although cDC2 are conventionally linked to Th2 and Th17 (autoimmunity) immunomodulation, cDC2 have been reported to have the potential to differentiate into inflammatory DCs, which function in a manner similar to cDC1 [47]. Hence, we postulate that the loss of both cDC1 and cDC2 would have hindered the development and maintenance of an adaptive response for protection against virulent ASFV. In this study, we only investigated the changes in cDC dynamics in early ASFV infection; there is a need to investigate the effects of virulent ASFV on the function of cDCs further as these cells have also been implicated in crosstalk with γδ-TCR^+^ cells [48].

Although both macrophages and monocytes are target cells for ASFV replication [31], macrophages have been found to be more susceptible to ASFV infection than monocytes *in vitro* [49]. CD163+ macrophages were broadly depleted across the tissues most likely due to replication of ASFV in these cells early in the infection course. While monocytes have been shown to infiltrate into tissues and replace the loss of macrophages after infection [32], clear expansion of monocytes was only observed in the spleen and SMLN and were reduced in the SMLN at 5 dpi. Mo-macrophages were also depleted by 5 dpi. Increase in monocytes within the spleen was previously observed using scRNAseq and from the proportions of infected monocytes detected in animals with the highest spleen viral loads, it is possible that the monocytes were infected after the macrophages and mo-macrophages were depleted [12]. Furthermore, monocytes have been shown to influence the upregulation of effector T cells in response to type I inflammation within LNs [50]. Hence, it is tempting to speculate that the reduction of monocytes in SMLN 5 dpi may have contributed to the lower levels of effector CTLs expressing perforin, but this remains to be explored in further detail.

While our p72 detection with antibodies was not as sensitive as qPCR or scRNAseq, the differential infection profiles observed in our inbred model demonstrates the diversity of cellular tropism of ASFV within the spleen, especially at high titres. In contrast to Zhu *et al.* [12] our study identified one animal (AZ48) that had a higher proportion of CD11b^−^ CADM1^+^CD172a^+^ DC2 cells that were infected in comparison to the other animals with advanced disease. Although we detected ASFV p72 in lymphocytes, our data (Supplementary Figure 38) is similar to previous data where between 0.12 – 0.26% of CD3+ and B cells were ASFV^+^ and these cells were shown to be non-permissive to ASFV infection [12].

## 6 Conclusion

In summary, our findings demonstrate a depletion of adaptive immune cells within the lymphocyte compartment, as well as a loss of professional APCs within the MPS. Direct infection and subsequent apoptosis are likely contributors to the depletion of MPS cells. This reduction of critical cell subsets, such as CD4+ T cells and cDCs, from both the innate and adaptive immune compartments in the early stages of ASFV infection further disrupts the bridge between these arms of immunity. Consequently, while there are initial attempts to initiate an adaptive immune response, this process is disrupted due to the absence of key immune cell populations required for its maintenance. The inability to generate a sufficiently robust or sustained adaptive immune response not only impairs immune control but may also contribute to accelerated disease progression and, ultimately, death. These findings highlight the need to investigate the innate-adaptive axis further with different ASFV isolates of varying virulence to determine if this immune imbalance is a defining feature of acute ASFV infection.

## 7 Acknowledgements

The authors would like to thank Jake Scales, Ollie Trussler, Zach Skoumbourdis, Louise Carder, Henry Steele, Dave Selby, Luke Fitzpatrick, Billy Matthews and Michael Collett for the care of the animals during the studies described in this manuscript and the flow cytometry facility for their support in this research.

## 8 Author contributions

Priscilla YL Tng (Conceptualisation, Data curation, Formal analysis, Funding acquisition, Investigation, Methodology, Supervision, Writing – original draft,, Writing – review & editing), Laila Al-Adwani (Methodology, Investigation, Writing – review & editing), Lynnette Goatley (Investigation, Writing – review & editing), Raquel Portugal (Investigation, Writing – review & editing), Anusyah Rathakrishnan (Investigation, Writing – review & editing), Christopher L Netherton (Conceptualisation, Funding acquisition, Resources, Supervision, Writing – review & editing)

## 11 Data availability

The data underlying this article are available on FigShare at https://dx.doi.org/10.6084/m9.figshare.c.7857272.

## 12 Ethical approval

All animal experiments were approved by the Animal Welfare and Ethical Review Board (AWERB) of The Pirbright institute and were conducted under the auspices of the Home Office Animals (Scientific Procedures) Act (ASPA, 1986). The animals were housed and cared for in accordance with the Code of Practice for the Housing and Care of Animals Bred, Supplied or Used for Scientific Purposes. To ensure high standards of animal welfare, bedding and species-specific enrichment were provided to the pigs throughout the study. Trained and qualified Personal License holders conducted all procedures under the oversight of Project License PPL70/8852. Throughout the studies, the pigs were under close supervision and were euthanised by an overdose of anaesthetic when they reached the scientific or humane endpoints.

